# An updated phylogeny of the *Alphaproteobacteria* reveals that the parasitic *Rickettsiales* and *Holosporales* have independent origins

**DOI:** 10.1101/462648

**Authors:** Sergio A. Muñoz-Gómez, Sebastian Hess, Gertraud Burger, B. Franz Lang, Edward Susko, Claudio H. Slamovits, Andrew J. Roger

## Abstract

The *Alphaproteobacteria* is an extraordinarily diverse and ancient group of bacteria. Previous attempts to infer its deep phylogeny have been plagued with methodological artefacts. To overcome this, we analyzed a dataset of 200 single-copy and conserved genes and employed diverse strategies to reduce compositional artefacts. Such strategies include using novel dataset-specific profile mixture models and recoding schemes, and removing sites, genes and taxa that are compositionally biased. We show that the *Rickettsiales* and *Holosporales* (both groups of intracellular parasites of eukaryotes) are not sisters to each other, but instead, the *Holosporales* has a derived position within the *Rhodospirillales*. Furthermore, we find that the *Rhodospirillales* might be paraphyletic and that the *Geminicoccaceae* could be sister to all ancestrally free-living alphaproteobacteria. Our robust phylogeny will serve as a framework for future studies that aim to place mitochondria, and novel environmental diversity, within the *Alphaproteobacteria*.

## INTRODUCTION

The *Alphaproteobacteria* is an extraordinarily diverse and disparate group of bacteria and well-known to most biologists for also encompassing the mitochondrial lineage (Williams, Sobral, and Dickerman 2007; Roger, Muñoz-Gómez, and Kamikawa 2017). The *Alphaproteobacteria* has massively diversified since its origin, giving rise to, for example, some of the most abundant (e.g., *Pelagibacter ubique*) and metabolically versatile (e.g., *Rhodobacter sphaeroides*) cells on Earth (Giovannoni 2017; Madigan, Jung, and Madigan 2009). The basic structure of the tree of the *Alphaproteobacteria* has largely been revealed through the analyses of 16S rRNA genes and several conserved proteins (Garrity 2005; Lee et al. 2005; Rosenberg et al. 2014; Fitzpatrick, Creevey, and McInerney 2006; Williams, Sobral, and Dickerman 2007; Brindefalk et al. 2011; Georgiades et al. 2011; Thrash et al. 2011; Luo 2015). Today, eight major orders are well recognized, namely the *Caulobacterales*, *Rhizobiales*, *Rhodobacterales*, *Pelagibacterales*, *Sphingomonadales*, *Rhodospirillales, Holosporales* and *Rickettsiales* (the latter two formerly grouped into the *Rickettsiales sensu lato*), and their interrelationships have also recently become better understood (Viklund, Ettema, and Andersson 2012; Viklund et al. 2013; Rodríguez-Ezpeleta and Embley 2012; Wang and Wu 2014, 2015). These eight orders were grouped into two subclasses by Ferla *et al*. (2013): the subclass *Rickettsiidae* comprising the order *Rickettsiales* and *Pelagibacterales*, and the subclass *Caulobacteridae* comprising all other orders.

The great diversity of the *Alphaproteobacteria* itself presents a challenge to deciphering the deepest divergences within the group. Such diversity encompasses a broad spectrum of genome (nucleotide) and proteome (amino acid) compositions (e.g., the A+T%-rich *Pelagibacterales versus* the G+C%-rich *Acetobacteraceae*) and molecular evolutionary rates (e.g., the fast-evolving *Pelagibacteriales*, *Rickettsiales* or *Holosporales versus* many slow-evolving species in the *Rhodospirillales*) (Ettema and Andersson 2009). This diversity may lead to pervasive artefacts when inferring the phylogeny of the *Alphaproteobacteria*, e.g., long-branch attraction (LBA) between the *Rickettsiales* and *Pelagibacterales*, especially when including mitochondria (Rodríguez-Ezpeleta and Embley 2012; Viklund, Ettema, and Andersson 2012; Viklund et al. 2013).

Moreover, there are still important unknowns about the deep phylogeny of the *Alphaproteobacteria* (Williams, Sobral, and Dickerman 2007; Ferla et al. 2013), for example, the divergence order among the *Rhizobiales*, *Rhodobacterales* and *Caulobacterales* (Williams, Sobral, and Dickerman 2007), the monophyly of the *Pelagibacterales* (Viklund et al. 2013) and the *Rhodospirillales* (Ferla et al. 2013), and the precise placement of the *Rickettsiales* and its relationship to the *Holosporales* (Wang and Wu 2015; Martijn et al. 2018).

Systematic errors stemming from using over-simplified evolutionary models are perhaps the major confounding and limiting factor to inferring deep evolutionary relationships; the number of taxa and genes (or sites) can also be important factors. Previous multi-gene tree studies of the *Alphaproteobacteria* were compromised by at least one of these problems, namely, simple or unrealistic evolutionary models (because they were not available at the time; e.g., Williams, Sobral, and Dickerman 2007), poor taxon sampling (because the focus was too narrow or few genomes were available; e.g., Williams, Sobral, and Dickerman 2007; Georgiades et al. 2011; Martijn et al. 2015) or a small number of genes (because the focus was mitochondria; e.g., Rodríguez-Ezpeleta and Embley 2012; Wang and Wu 2015; Martijn et al. 2018). The most recent study on the phylogeny of the *Alphaproteobacteria*, and mitochondria, attempted to counter systematic errors (or phylogenetic artefacts) by reducing amino acid compositional heterogeneity (Martijn et al. 2018). Even though some deep relationships were not robustly resolved, these analyses suggested that the *Pelagibacterales*, *Rickettsiales* and *Holosporales*, which have compositionally biased genomes, are not each other’s closest relatives (Martijn et al. 2018). A resolved and robust phylogeny of the *Alphaproteobacteria* is fundamental to addressing questions such as how streamlined bacteria, intracellular parasitic bacteria, or mitochondria evolved from their alphaproteobacterial ancestors. Therefore, a systematic study of the different biases affecting the phylogeny of the *Alphaproteobacteria*, and its underlying data, is much needed.

Here, we revised the phylogeny of the *Alphaproteobacteria* by using a large dataset of 200 conserved single-copy genes and employing carefully designed strategies aimed at alleviating phylogenetic artefacts. We found that amino acid compositional heterogeneity, and more generally long-branch attraction, were major confounding factors in estimating phylogenies of the *Alphaproteobacteria*. In order to counter these biases, we used novel dataset-specific profile mixture models and recoding schemes (both specifically designed to ameliorate compositional heterogeneity), and removed sites, genes and taxa that were compositionally biased. We also present three draft genomes for endosymbiotic alphaproteobacteria belonging to the *Rickettsiales* and *Holosporales*: (1) an undescribed midichloriacean endosymbiont of *Peranema trichophorum*, (2) an undescribed rickettsiacean endosymbiont of *Stachyamoeba lipophora*, and (3) the holosporalean ‘*Candidatus* Finniella inopinata’, an endosymbiont of the rhizarian amoeboflagellate *Viridiraptor invadens* (Hess, Suthaus, and Melkonian 2015). Our results provide the first strong evidence that the *Holosporales* are unrelated to the *Rickettsiales* and originated instead from within the *Rhodospirillales*. We incorporate these and other insights regarding the deep phylogeny of the *Alphaproteobacteria* into an updated taxonomy.

## RESULTS

### The genomes and phylogenetic positions of three novel endosymbiotic alphaproteobacteria (Rickettsiales and Holosporales)

We sequenced the genomes of the novel holosporalean ‘*Candidatus* Finniella inopinata’, an endosymbiont of the rhizarian amoeboflagellate *Viridiraptor invadens* (Hess, Suthaus, and Melkonian 2015), and two undescribed rickettsialeans, one associated with the heterolobosean amoeba *Stachyamoeba lipophora* and the other with the euglenoid flagellate *Peranema trichophorum*. The three genomes are small with a reduced gene number and high A+T% content, strongly suggesting an endosymbiotic lifestyle (Table 1). Comparisons of their rRNA genes show that these genomes are truly novel, being considerably divergent from other described alphaproteobacteria. As of February 2018, the closest 16S rRNA gene to that of the *Stachyamoeba*-associated rickettsialean belongs to *Rickettsia massiliae* str. AZT80, with only 88% identity. On the other hand, the closest 16S rRNA gene to that of the *Peranema*-associated rickettsialean belongs to an endosymbiont of *Acanthamoeba* sp. UWC8, which is only 92% identical. Phylogenetic analysis of both the 16S rRNA gene and the 200-gene set confirm that each species belongs to different families and orders within the *Alphaproteobacteria* (Fig. S4). ‘*Candidatus* Finniella inopinata’ belongs to the recently described ‘*Candidatus* Paracaedibacteraceae’ in the *Holosporales* (Hess, Suthaus, and Melkonian 2015), whereas the *Stachyamoeba*-associated rickettsialean belongs to the *Rickettsiaceae*, and the *Peranema*-associated rickettsialean belongs to the ‘*Candidatus* Midichloriaceae’, in the *Rickettsiales*.

**Table 1.** Genome features for the three novel rickettsialeans sequenced in this study.

| Species | '*Candidatus* Finniella inopinata' | *Stachyamoeba*-associated rickettsialean | *Peranema*-associated rickettsialean |
| --- | --- | --- | --- |
| Genome size | 1,792,168 bp | 1,738,386 bp | 1,375,759 bp |
| N50 | 174,737 bp | 1,738,386 bp | 28,559 bp |
| Contig number | 28 | 1 | 125 |
| Gene number<sup>1</sup> | 1,741 | 1,588 | 1,223 |
| A+T% content | 56.58% | 67.01% | 59.13% |
| Family | *Paracaedibacteraceae* | *Rickettsiaceae* | '*Candidatus* Midicloriaceae' |
| Order | *Holosporales* | *Rickettsiales* | *Rickettsiales* |
| Completeness<sup>2</sup> | 94.96% | 97.12% (=100%) | 92.08% |
| Redundancy<sup>2</sup> | 0.0% | 0.0% | 2.1% |

<sup>1</sup> as predicted by prokka v.1.13 (rRNA genes were searched with BLAST).
<sup>2</sup> as estimated by anvi'o v.2.4.0 using the Campbell et al., 2013 marker set.

### Compositional heterogeneity appears to be a major confounding factor affecting phylogenetic inference of the Alphaproteobacteria

The average-linkage clustering of amino acid compositions shows that the *Rickettsiales*, *Pelagibacterales* and *Holosporales* are clearly distinct from other alphaproteobacteria. This indicates that these three taxa have divergent proteome amino acid compositions (Fig. 1A). These taxa also have the lowest GARP:FIMNKY ratios in all the *Alphaproteobacteria* (Fig. 1A); the *Pelagibacterales* being the most divergent, followed by the *Rickettsiales* and then the *Holosporales*. Such biased amino acid compositions appear to be the consequence of genome nucleotide compositions that are strongly biased towards high A+T%—a scatter plot of genome G+C% and proteome GARP:FIMNKY ratios shows a similar clustering of the *Rickettsiales*, *Pelagibacterales* and *Holosporales* (Fig. 1B). This compositional similarity in the proteomes of the *Rickettsiales*, *Pelagibacterales* and *Holosporales*, which also turn out to be the longest-branching alphaproteobacterial groups in previously published phylogenies (e.g., Wang and Wu 2015), could be the outcome of either a shared evolutionary history (i.e., the groups are most closely related to one another), or alternatively, evolutionary convergence (e.g., because of similar lifestyles or evolutionary trends toward small cell and genome sizes).

**Figure 1.**
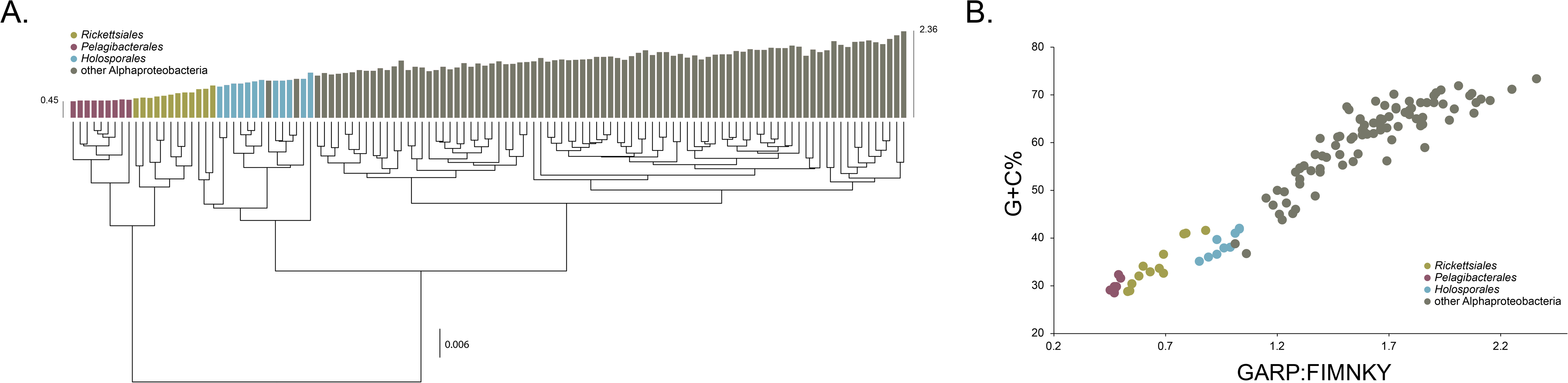
Compositional heterogeneity in the *Alphaproteobacteria* is a major factor that confounds phylogenetic inference. There are great disparities in the genome G+C% content and amino acid compositions of the *Rickettsiales*, *Pelagibacterales* and *Holosporales* with all other alphaproteobacteria. **A.** A UPGMA (average-linkage) clustering of amino acid compositions (based on the 200 gene set for the *Alphaproteobacteria*) shows that the *Rickettsiales* (brown), *Pelagibacterales* (gray), and *Holosporales* (light blue) all have very similar proteome amino acid compositions. At the tips of the tree, GARP:FIMNKY ratio values are shown as bars. **B.** A scatterplot depicting the strong correlation between G+C% (nucleotide compositions) and GARP:FIMNKY ratios (amino acid composition) for the 120 taxa in the *Alphaproteobacteria* shows a similar clustering of the *Rickettsiales*, *Pelagibacterales* and *Holosporales*.

As a first step to discriminate between these two alternatives, we used maximum likelihood to estimate a tree on our 200-gene dataset for the *Alphaproteobacteria* under the site-heterogenous model LG+PMSF(ES60)+F+R6. The resulting tree united the *Rickettsiales*, *Pelagibacterales* and *Holosporales* in a fully supported clade (Fig. 2A). The clustering of these three groups is suggestive of a phylogenetic artefact (e.g., long-branch attraction or LBA); indeed, such a pattern resembles the one seen in the tree of proteome amino acid compositions (see Fig. 1A). This is because the three groups have the longest branches in the *Alphaproteobacteria* tree and have compositionally biased and fast-evolving genomes (see Fig. 2). If evolutionary convergence in amino acid compositions is confounding phylogenetic inference for the *Alphaproteobacteria*, methods aimed at reducing compositional heterogeneity might disrupt the clustering of the *Rickettsiales*, *Pelagibacterales* and *Holosporales*.

**Figure 2.**
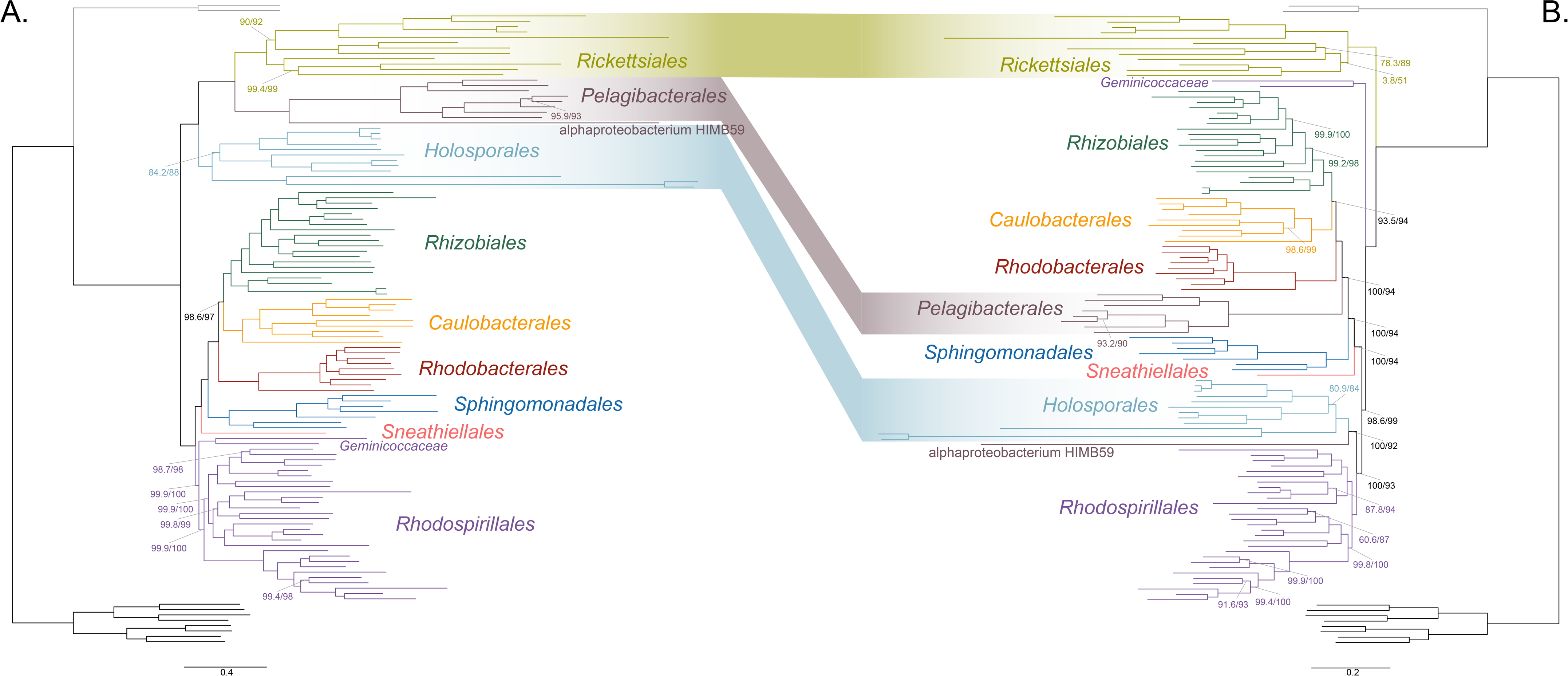
Decreasing compositional heterogeneity by removing compositionally biased sites disrupts the clustering of the *Rickettsiales*, *Pelagibacterales* and *Holosporales*. All branch support values are 100% SH-aLRT and 100% UFBoot unless annotated. **A.** A maximum-likelihood tree inferred under the LG+PMSF(ES60)+F+R6 model and from the intact dataset which is highly compositionally heterogeneous. The three long-branching orders, the *Rickettsiales Pelagibacterales* and *Holosporales*, that have similar amino acid compositions form a clade. **B.** A maximum-likelihood tree inferred under the LG+PMSF(ES60)+F+R6 model and from a dataset whose compositional heterogeneity has been decreased by removing 50% of the most biased sites according to 𝒵. In this phylogeny the clustering of the *Rickettsiales*, *Pelagibacterales* and *Holosporales* is disrupted. The *Pelagibacterales* is sister to the *Rhodobacterales*, *Caulobacterales* and *Rhizobiales*. The *Holosporales* becomes sister to the *Rhodospirillales*. The *Rickettsiales* retains its basal position as sister to the *Caulobacteridae*. See Fig. S6 for the Bayesian consensus trees inferred in PhyloBayes MPI v1.7 under the CAT-Poisson+Γ4 model.

To further test whether the clustering of the *Rickettsiales*, *Pelagibacterales* and *Holosporales* is real or artefactual we used several different strategies to reduce the compositional heterogeneity of our dataset (see Fig. S2 for the diverse strategies employed). When removing the 50% most compositionally biased (heterogeneous) sites according to 𝒵, the clustering between the *Rickettsiales*, *Pelagibacterales* and *Holosporales* is disrupted (Fig. 2B). The new more derived placements for the *Pelagibacterales* and *Holosporales* are well supported (further described below), and support tends to increase as compositionally biased sites are removed (Fig. S8). Furthermore, when each of these long-branching taxa is analyzed in isolation (i.e., in the absence of the other two), and compositionally heterogeneity is decreased, new phylogenetic patterns emerge that are incompatible, or in conflict, with their clustering (Fig. S10-S13). In other words, removing the most compositionally biased sites, recoding the data into reduced character-state alphabets to minimize compositional bias, or using only the most compositionally homogeneous genes, converge to very similar phylogenetic patterns for the *Alphaproteobacteria* that are however incompatible with the clustering of the *Rickettsiales*, *Pelagibacterales* and *Holosporales* (e.g., Fig. S10-S13, and 3). On the other hand, removing fast-evolving sites does not disrupt the clustering of the these three long-branching groups (Fig. S7), suggesting that high evolutionary rates per site are not a major confounding factor when inferring the phylogeny of the *Alphaproteobacteria*.

### The Holosporales is unrelated to the Rickettsiales and is instead most likely derived within the Rhodospirillales

The *Holosporales* has traditionally been considered part of the *Rickettsiales sensu lato* because it appears as sister to the *Rickettsiales* in many trees. It is exclusively composed of endosymbiotic bacteria living within diverse eukaryotes, and such a lifestyle is shared with all other members of the *Rickettsiales*. When we decrease, and then account for compositional heterogeneity, we recover tree topologies in which the *Holosporales* moves away from the *Rickettsiales* (e.g., Fig. 2B and S13A). For example, the *Holosporales* becomes sister to all free-living alphaproteobacteria (the *Caulobacteridae*) when only the 40 most homogeneous genes are used (Fig. S13A) or when 10% of the most compositionally biased sites are removed (Fig. S8). When compositional heterogeneity is further decreased by removing 50% of the most compositionally biased sites, the *Holosporales* becomes sister to the *Rhodospirillales* (Fig. 2B and S8; and see also Fig. S10B and S13B).

Similarly, when the long-branching *Rickettsiales* and *Pelagibacterales* groups (plus the extremely long-branching genera *Holospora* and ‘*Candidatus* Hepatobacter’) are removed, after compositional heterogeneity had been decreased through site removal, the *Holosporales* move to a much more derived position well within the *Rhodospirillales* (Fig. 3A). If the very compositionally biased and fast-evolving *Holospora* and ‘*Candidatus* Hepatobacter’ are left in, the *Holosporales* are pulled away from its derived position and the whole clade moves closer to the base of the tree (data not shown). The same behaviour is seen when these same taxa are removed, and the data are then recoded into four- or six-character states (Fig. 3B and S14). Specifically, the *Holosporales* now consistently branches as sister to a subgroup of rhodospirillaleans that includes, among others, the epibiotic predator *Micavibrio aeruginosavorus* and the purple nonsulfur bacterium *Rhodocista centenaria* (the *Azospirillaceae*, see below). This new placement of the *Holosporales* has nearly full support under both maximum likelihood and Bayesian inference (e.g., >95% UFBoot; see Fig. 3).Thus, three different analyses independently converge to the same pattern and support a derived origin of the *Holosporales* within the *Rhodospirillales*: (1) removal of compositionally biased sites (Fig. 3A), (2) data recoding into four-character states using the dataset-specific scheme S4 (Fig. 3B), and (3) data recoding into six-character states using the dataset-specific scheme S6 (Fig. S14); each of these strategies had to be combined with the removal of the *Pelagibacterales* and *Rickettsiales* to recover this phylogenetic position for the *Holosporales*.

**Figure 3.**
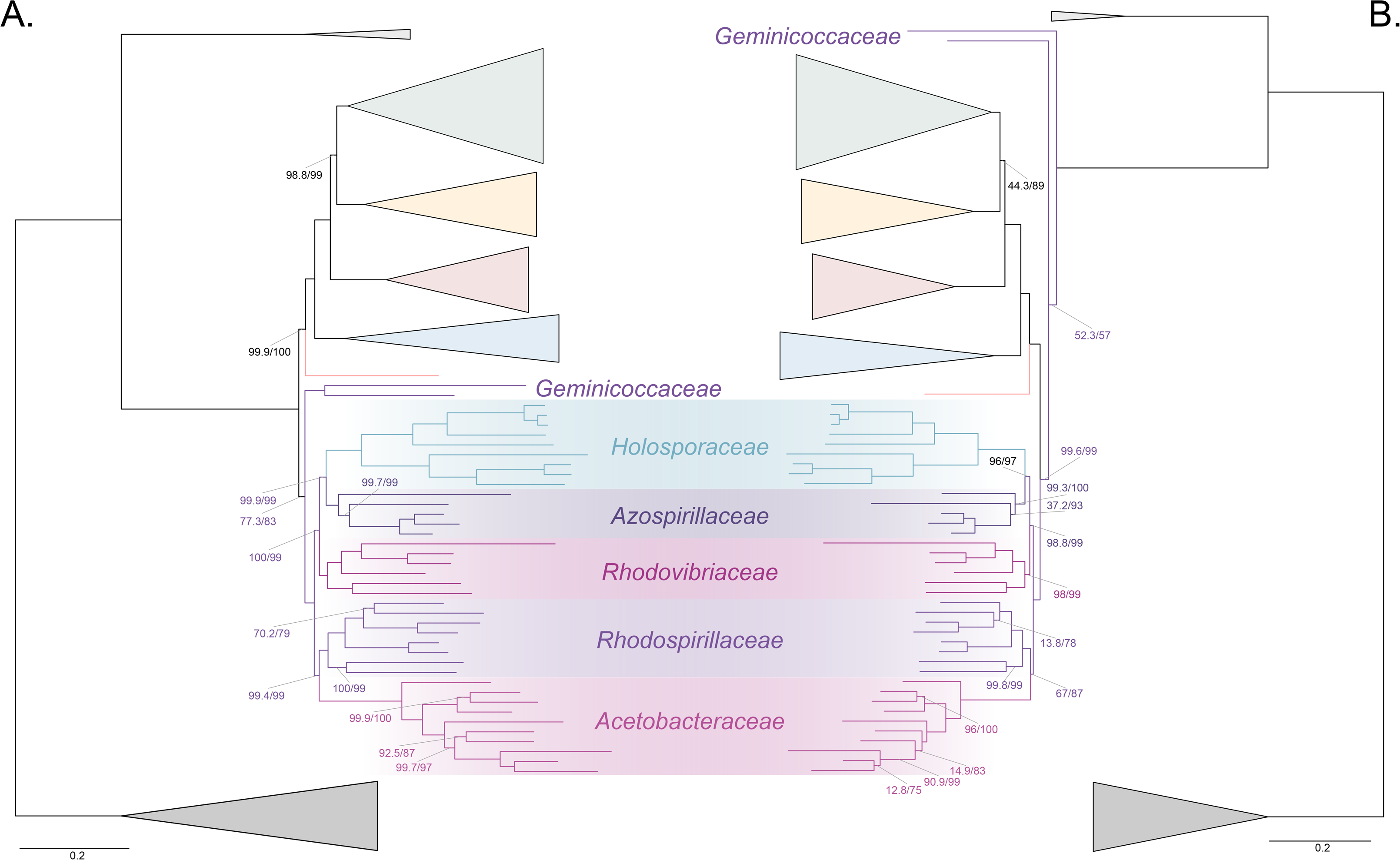
The *Holosporales* branches in a derived position within the *Rhodospirillales* when compositional heterogeneity is reduced and the long-branching *Rickettsiales* and *Pelagibacterales* are removed. Branch support values are 100% SH-aLRT and 100% UFBoot unless annotated. **A.** A maximum-likelihood tree, inferred under the LG+PMSF(ES60)+F+R6 model, to place the *Holosporales* in the absence of the *Rickettsiales* and the *Pelagibacterales* and when compositional heterogeneity has been decreased by removing 50% of the most biased sites. The *Holosporales* is sister to the *Azospirillaceae* fam. nov. within the *Rhodospirillales*. **B.** A maximum-likelihood tree, inferred under the GTR+ES60S4+F+R6 model, to place the *Holosporales* in the absence of the *Rickettsiales* and the *Pelagibacterales* and when the data have been recoded into a four-character state alphabet (the dataset-specific recoding scheme S4: ARNDQEILKSTV GHY CMFP W) to reduce compositional heterogeneity. This phylogeny shows a pattern that matches that inferred when compositional heterogeneity has been alleviated through site removal. See Fig. S9 for the Bayesian consensus trees inferred in PhyloBayes MPI v1.7 and under the and the CAT-Poisson+Γ4 model.

A fourth independent analysis further supports a derived placement of the *Holosporales* nested within the *Rhodospirillales*. Bayesian inference using the CAT-Poisson+Γ4 model, on a dataset whose compositional heterogeneity had been decreased by removing 50% of the most compositionally biased sites but for which no taxa had been removed, also recovered the *Holosporales* as sister to the *Azospirillaceae* (see Fig. S6).

### The Rhodospirillales is a diverse order and comprises five well-supported families

The *Rhodospirillales* is an ancient and highly diversified group, but unfortunately this is rarely obvious from published phylogenies because most studies only include a few species for this order (Williams, Sobral, and Dickerman 2007; Georgiades et al. 2011; Ferla et al. 2013). We have included a total of 31 *Rhodospirillales* taxa to better cover its diversity. Such broad sampling reveals trees with five clear subgroups within the *Rhodospirillales* that are well-supported in most of our analyses (e.g., Fig. 2B and 3). First is the *Acetobacteraceae* which comprises acetic acid, acidophilic, and photosynthesizing (bacteriochlorophyll-containing) bacteria. The *Acetobacteraceae* is strongly supported and relatively divergent from all other families within the *Rhodospirillales*. Sister to the *Acetobacteraceae* is another subgroup that comprises many photosynthesizing bacteria, including the type species for the *Rhodospirillales*, *Rhodospirillum rubrum*, as well as the magnetotactic bacterial genera *Magnetospirillum*, *Magnetovibrio* and *Magnetospira* (Fig. 3). This subgroup best corresponds to the poorly defined and paraphyletic *Rhodospirillaceae* family. We amend the *Rhodospirillaceae* taxon and restrict it to the clade most closely related to the *Acetobacteraceae*. As described above, when artefacts are accounted for, the *Holosporales* most likely branches within the *Rhodospirillales* and therefore we suggest the *Holosporales sensu* Szokoli *et al*. (2016) is lowered in rank to the family *Holosporaceae*, which is sister to the *Azospirillaceae* (Fig. 3). The *Azospirillaceae* **fam. nov.** contains the purple bacterium *Rhodocista centenaria* and the epibiotic (neither periplasmic nor intracellular) predator *Micavibrio aeruginosavorus*, among others. The *Holosporaceae* and the *Azospirillaceae* clades appear to be sister to the *Rhodovibriaceae* **fam. nov.** (Fig. 3), a well-supported group that comprises the purple nonsulfur bacterium *Rhodovibrio salinarum*, the aerobic heterotroph *Kiloniella laminariae*, and the marine bacterioplankter ‘*Candidatus* Puniceispirillum marinum’ (or the SAR116 clade). Each of these subgroups and their interrelationships—with the exception of the *Holosporaceae* that branches within the *Rhodospirillales* only after compositional heterogeneity is countered—are strongly supported in nearly all of our analyses (e.g., see Fig. 2B and 3).

### The Geminicoccaceae might be basal to all other free-living alphaproteobacteria (the Caulobacteridae)

The *Geminicocacceae* is a recently proposed family within the *Rhodospirillales* (Proença et al. 2017). It is currently represented by only two genera, *Geminicoccus* and *Arboriscoccus* (Foesel et al. 2007; Proença et al. 2017). In most of our trees, however, *Tistrella mobilis* is often sister to *Geminicoccus roseus* with full statistical support (e.g., Fig. 2B and 3A, but see Fig. S9 for an exception) and we therefore consider it to be part of the *Geminococcaceae*. Interestingly, the *Geminicoccaceae* tends to have two alternative stable positions in our analyses, either as sister to all other families of the *Rhodospirillales* (e.g., Fig. 2A and 3A), or alternatively, as sister to all other orders of the *Caulobacteridae* (i.e., representing the most basal lineage of free-living alphaproteobacteria; Fig. 2B and 3B, or Fig. S11C and S12C). Our analyses designed to alleviate compositional heterogeneity, specifically site removal and recoding (without taxon removal), favor the latter position for the *Geminicoccaceae* (Fig. 2B and 3B). Moreover, as compositionally biased sites are progressively removed, support for the affiliation of the *Geminicoccaceae* with the *Rhodospirillales* decreases, and after 50% of the sites have been removed, the *Geminicoccaceae* emerges as sister to all other free-living alphaproteobacteria with strong support (>95% UFBoot; Fig. S8). In further agreement with this trend, the much simpler model LG4X places the *Geminicocacceae* in a derived position as sister to the *Acetobacteraceae* (data not shown), but as model complexity increases, and compositional heterogeneity is reduced, the *Geminicoccaceae* moves closer to the base of the *Alphaproteobacteria* (Fig. 2A and 3A). Such a placement suggests that the *Geminicoccaceae* may be a novel and independent order-level lineage in the *Alphaproteobacteria*. However, because of the uncertainty in our results we opt here for conservatively keeping the *Geminicoccaceae* as the sixth family of the *Rhodospirillales* (Fig. 3A).

### Other deep relationships in the Alphaproteobacteria (Pelagibacterales, Rickettsiales, alphaproteobacterium sp. HIMB59)

The clustering of the *Pelagibacterales* (formerly the SAR11 clade) with the *Rickettsiales* and *Holosporales* is more easily disrupted than that of the *Holosporales*. The removal of compositionally biased sites (from 30% on; 16,320 out of 54,400 sites; Fig. S8), data recoding into four-character states (Fig. S12D), and a set of the most compositionally homogeneous genes (Fig. S13D), all support a derived placement of the *Pelagibacterales* as sister to the *Rhodobacterales*, *Caulobacterales* and *Rhizobiales* (in agreement with Rodríguez-Ezpeleta and Embley 2012; Viklund, Ettema, and Andersson 2012; Viklund et al. 2013; Martijn et al. 2018). The *Caulobacterales* is sister to the *Rhizobiales*, and the *Rhodobacterales* sister to both (e.g., Fig. 2B and 3). This is consistent throughout most of our results and such interrelationships become very robustly supported as compositional heterogeneity is increasingly alleviated (Fig. S8). The placement of the *Rickettsiales* as sister to the *Caulobacteridae* (i.e., all other alphaproteobacteria) remains stable across different analyses (Fig. 2B, S10C, S11C, S12C and S13D); this is also true when the other long-branching taxa, the *Pelagibacterales* and *Holosporales*, are removed. Yet, the interrelationships inside the *Rickettsiales* order remain uncertain; the ‘*Candidatus* Midichloriaceae’ becomes sister to the *Anaplasmataceae* when fast sites are removed (Fig. S7), but to the *Rickettsiaceae* when compositionally biased sites are removed (Fig. S8). The placement of alphaproteobacterium sp. HIMB59 is entirely uncertain (e.g., see Fig. 2B and S11E; in contrast to Grote et al. 2012); it does not have a stable position in the *Alphaproteobacteria* across our diverse analyses. This is consistent with previous reports that suggest that alphaproteobacterium sp. HIMB59 is not closely related to the *Pelagibacterales* (Viklund et al. 2013; Martijn et al. 2018).

## DISCUSSION

We have employed a diverse set of strategies to investigate the phylogenetic signal contained within 200 genes for the *Alphaproteobacteria*. Specifically, such strategies were primarily aimed at reducing amino acid compositional heterogeneity among taxa— a phenomenon that permeates our dataset (Fig. 1). Compositional heterogeneity is a clear violation of the phylogenetic models used in our, and previous, analyses, and known to cause phylogenetic artefacts (Foster 2004). In the absence of more sophisticated models for inferring deep phylogeny, the only way to counter artefacts caused by compositional heterogeneity is by removing compositionally biased sites or taxa, or recoding amino acids into reduced alphabets. A combination of these strategies reveals that the *Rickettsiales sensu lato* (i.e., the *Rickettsiales* and *Holosporales*) is polyphyletic. Our analyses suggest that the *Holosporales* is derived within the *Rhodospirillales*, and that therefore this taxon should be lowered in rank and renamed the *Holosporaceae* family (see Fig. 2B and 3). The same methods suggest that the *Rhodospirillales* might indeed be a paraphyletic order and that the *Geminicoccaceae* could be a separate lineage that is sister to the *Caulobacteridae* (e.g., Fig. 2B). These two results, combined with our broader sampling, reorganize the internal phylogenetic structure of the *Rhodospirillales* and show that its diversity can be grouped into at least five well-supported major families (Fig. 3).

In 16S rRNA gene trees, the *Holosporales* has most often been allied to the *Rickettsiales* (Montagna et al. 2013; Hess, Suthaus, and Melkonian 2015). The apparent diversity of this group has quickly increased in recent years as more and more intracellular bacteria living within protists have been described (e.g., Hess, Suthaus, and Melkonian 2015; Szokoli et al. 2016; Eschbach et al. 2009; Boscaro et al. 2013). An endosymbiotic lifestyle is shared by all members of the *Holosporales* and is also shared with all those that belong to the *Rickettsiales*. Thus, it had been reasonable to accept their shared ancestry as suggested by some 16S rRNA gene trees (e.g., Montagna et al. 2013; Santos and Massard 2014; Hess, Suthaus, and Melkonian 2015). Apparent strong support for the monophyly of the *Rickettsiales* and the *Holosporales* recently came from some multi-gene trees by Wang and Wu (2014, 2015) who expanded sampling for the *Holosporales*. However, an alternative placement for the *Holosporales* as sister to the *Caulobacteridae* has been reported by Ferla et al. (2013) based on rRNA genes, by Georgiades et al., (2011) based on 65 genes, by Schulz et al., (2015) based on 139 genes, as well as by Wang and Wu (2015) based on 26, 29, or 200 genes (see the supplementary information in Wang and Wu, 2015). This placement was acknowledged by Szokoli et al. (2015), who formally established the order *Holosporales*. Most recently, Martijn et al. 2018, who used strategies to reduce compositional heterogeneity, and similarly to Wang and Wu (2015), recovered a number of placements for the *Holosporales* within the *Alphaproteobacteria*; however these different placements for the *Holosporales* were poorly supported. Here we provide strong evidence for the hypothesis that the *Holosporales* is not related to the *Rickettsiales*, as suggested earlier (Georgiades et al. 2011; Ferla et al. 2013; Szokoli et al. 2016). The *Rickettsiales sensu lato* is polyphyletic. We show that the *Holosporales* is artefactually attracted to the *Rickettsiales* (e.g., Fig. 2A), but as compositional bias is increasingly alleviated (through site removal and recoding), they move further away from them (Fig. 2B). The *Holosporales* is placed within the *Rhodopirillales* as sister to the family *Azospirillaceae* (Fig. 3). The similar lifestyles of the *Holosporales* and *Rickettsiales*, as well as other features like the presence of an ATP/ADP translocase (Wang and Wu 2014), are therefore likely the outcome of convergent evolution.

A derived origin of the *Holosporales* has important implications for understanding the origin of mitochondria and the nature of their ancestor. Wang and Wu (2014, 2015) proposed that mitochondria are phylogenetically embedded within the *Rickettsiales sensu lato*. In their trees, mitochondria were sister to a clade formed by the *Rickettsiaceae*, *Anaplasmataceae* and ‘*Candidatus* Midichloriaceae’, and the *Holosporales* was itself sister to all of them. This phylogenetic placement for mitochondria suggested that the ancestor of mitochondria was an intracellular parasite (Wang and Wu, 2014). But if the *Holosporales* is a derived group of rhodospirillaleans as we show here (see Fig. 3), then the argument that mitochondria necessarily evolved from parasitic alphaproteobacteria no longer holds. While the sisterhood of mitochondria and the *Rickettsiales sensu stricto* is still a possibility, such a relationship does not imply that the two groups shared a parasitic common ancestor (i.e., a parasitic ancestry for mitochondria). The most recent analyses done by Martijn *et al*. (2018) suggest that mitochondria are sister to all known alphaproteobacteria, also suggesting their non-parasitic ancestry. Our study, and that of Martijn et al., thus complement each other and support the view that mitochondria most likely evolved from ancestral free-living alphaproteobacteria (*contra* Sassera et al. 2011; Wang and Wu 2014, 2015).

The order *Rhodospirillales* is quite diverse and includes many purple nonsulfur bacteria as well as all magnetotactic bacteria within the *Alphaproteobacteria*. The *Rhodospirillales* is sister to all other orders in the *Caulobacteridae*, and has historically been subdivided into two families: the *Rhodospirillaceae* and the *Acetobacteraceae*. Recently, a new family, the *Geminicoccaceae*, was established for the *Rhodospirillales* (Proença et al. 2017). However, some of our analyses suggest that the *Geminicoccaceae* might be sister to all other *Caulobacteridae* (e.g., Fig. 2B and 3B). This phylogenetic pattern, therefore, suggests that the *Rhodospirillales* may be a paraphyletic order. The placement of the *Geminicoccaceae* as sister to the *Caulobacteridae* needs to be further tested once more sequenced diversity for this group becomes available; if it were to be confirmed, the *Geminicoccaceae* should be elevated to the order level. Whereas the *Acetobacteraceae* is phylogenetically well-defined, there has been considerable uncertainty about the *Rhodospirillaceae* (e.g., Ferla et al., 2013), primarily because of poor sampling and a lack of resolution provided by the 16S rRNA gene. We subdivide the *Rhodospirillaceae sensu lato* into three subgroups (Fig. 3). We restrict the *Rhodospirillaceae sensu stricto* to the subgroup that is sister to the *Acetobacteracae* (Fig. 3). The other two subgroups are the *Rhodovibriaceae* and the *Azospirillaceae*; the latter is sister to the *Holosporaceae* (Fig. 3).

Based on our fairly robust phylogenetic patterns, we have updated the higher-level taxonomy of the *Alphaproteobacteria* (Fig. 4). We exclude the *Magnetococcales* from the *Alphaproteobacteria* class because of its divergent nature (e.g., see Fig. 1 in Esser, Martin, and Dagan 2007 which shows that many of *Magnetococcus*’ genes are more similar to those of beta-, and gammaproteobacteria). In agreement with its intermediate phylogenetic placement, we endorse the *Magnetococcia* class as proposed by Parks et al. (2018). At the highest level we define the *Alphaproteobacteria* class as comprising two subclasses *sensu* Ferla et al. (2013), the *Rickettsidae* and the *Caulobacteridae*. The former contains the *Rickettsiales*, and the latter contains all other orders, which are primarily and ancestrally free-living alphaproteobacteria. The order *Rickettsiales* comprises three families as previously defined, the *Rickettsiaceae*, the *Anaplasmataceae*, and the ‘*Candidatus* Midichloriaceae’. On the other hand, the *Caulobacteridae* is composed of seven phylogenetically well-supported orders: the *Rhodospirillales*, *Sneathiellales*, *Sphingomonadales*, *Pelagibacterales*, *Rhodobacterales*, *Caulobacterales* and *Rhizobiales*. Among the many species claimed to represent new order-level lineages on the basis of 16S rRNA gene trees (Cho and Giovannoni 2003; Kwon et al. 2005; Kurahashi et al. 2008; Wiese et al. 2009; Harbison et al. 2017), only *Sneathiella* deserves order-level status (Kurahashi et al. 2008), since all others have derived placements in our trees and those published by others (Williams et al. 2012; Bazylinski et al. 2013; Venkata Ramana et al. 2013; Harbison et al. 2017). The *Rhodospirillales* order comprises six families, three of which are new, namely the *Holosporaceae*, *Azospirillaceae* and *Rhodovibriaceae* (Fig. 4). This new higher-level classification of the *Alphaproteobacteria* updates and expands those presented by Ferla et al. (2013), the ‘Bergey’s Manual of Systematics of Archaea and Bacteria’ (Garrity 2005; Whitman 2015), and ‘The Prokaryotes’ (Rosenberg et al. 2014). The classification scheme proposed here could be partly harmonized with that recently proposed by Parks et al., (2018) by elevating the six families within the *Rhodospirllales* to the order level; the phylograms by Parks et al., (2018), however, are in conflict with those shown here and many of their proposed taxa are as well.

**Figure 4.**
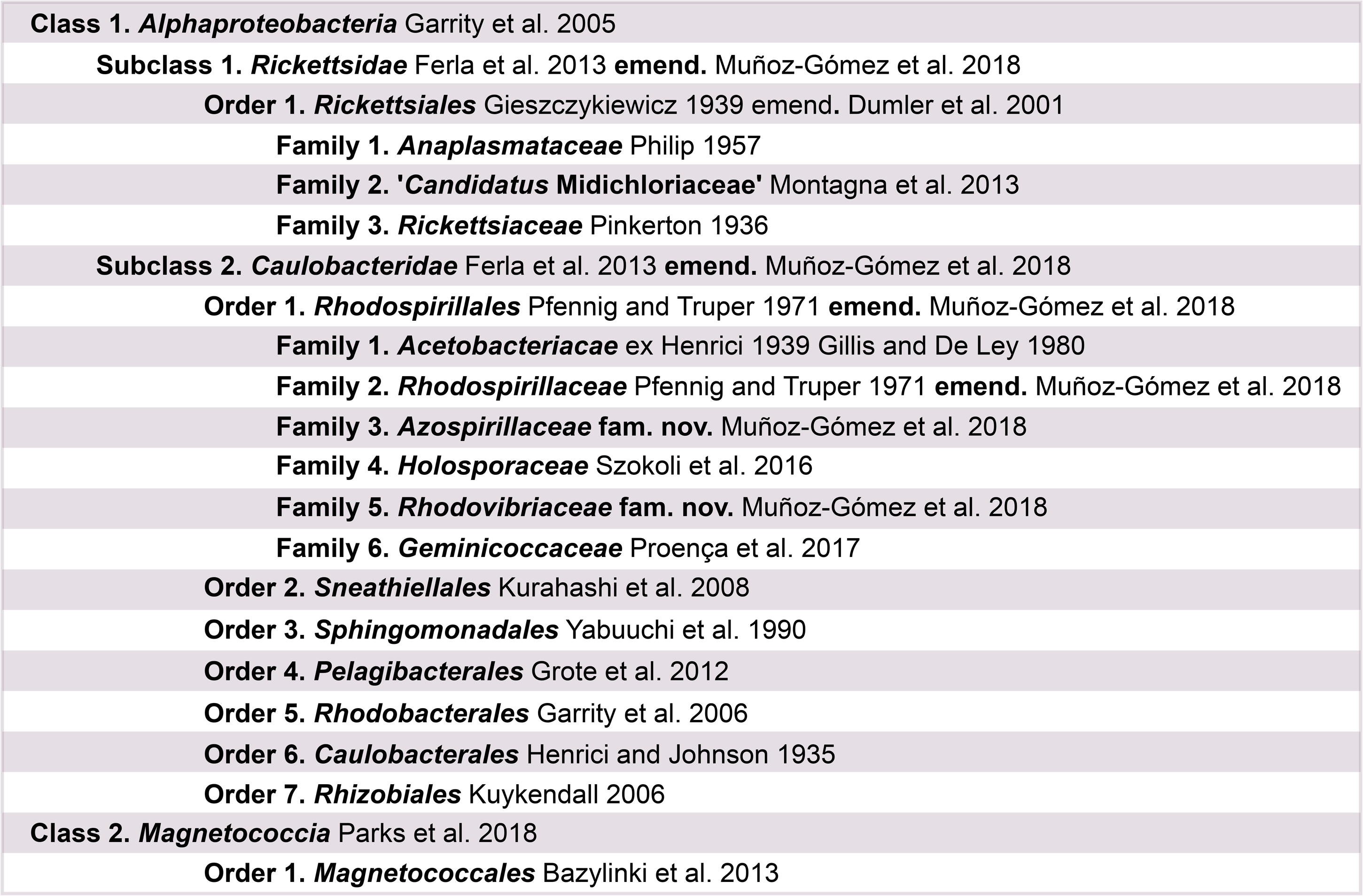
A higher-level classification scheme for the *Alphaproteobacteria* and the *Magnetococcia* classes within the *Proteobacteria*, and the *Rickettsiales* and *Rhodospirillales* orders within the *Alphaproteobacteria*.

## CONCLUSIONS

We employed a combination of methods to decrease compositional heterogeneity in order to disrupt artefacts that arise when inferring the phylogeny of the *Alphaproteobacteria*. This is an example of the complex nature of the historical signal contained in modern genomes and the limitations of our current evolutionary models to capture these signals. A robust phylogeny of the *Alphaproteobacteria* is a precondition for placing the mitochondrial lineage. This is because including mitochondria certainly exacerbates the already strong biases in the data, and therefore represents additional sources for artefact in phylogeny inference (as seen in Wang and Wu, 2015 where the *Holosporales* is attracted by both mitochondria and the *Rickettsiales*). Future endeavors should take into account the new phylogenetic framework developed in here. The incorporation of non-culturable ‘environmental’ diversity recovered from metagenomes will certainly expand the known diversity of the *Alphaproteobacteria* and improve phylogenetic inference.

## TAXON DESCRIPTIONS

*Rickettsidae* emend. (*Alphaproteobacteria*) *Rickettsia* is the type genus of the subclass. The *Rickettsidae* subclass is here amended by redefining its circumscription so it remains monophyletic by excluding the *Pelagibacterales* order. The emended *Rickettsidae* subclass within the *Alphaproteobacteria* class is defined based on phylogenetic analyses of 200 genes which are predominantly single-copy and vertically-inherited (unlikely laterally transferred) when compositionally heterogeneity was decreased by site removal or recoding. Phylogenetic (node-based) definition: the least inclusive clade containing *Anaplasma phagocytophilum* HZ, *Rickettsia typhi* Wilmington, and ‘*Candidatus* Midichloria mitochondrii’ IricVA. The *Rickettsidae* does not include: *Pelagibacter* sp. HIMB058, ‘*Candidatus* Pelagibacter sp.’ IMCC9063, alphaproteobacterium HIMB59, *Caedibacter* sp. 37-49, ‘*Candidatus* Nucleicultrix amoebiphila’ FS5, ‘*Candidatus* Finniella lucida’, *Holospora obtusa* F1, *Sneathiella glossodoripedis* JCM 23214, *Sphingomonas wittichii*, and *Brevundimonas subvibrioides* ATCC 15264.

*Caulobacteridae* emend. (*Alphaproteobacteria*) *Caulobacter* is the type genus of the subclass. The *Caulobacteridae* subclass is here amended by redefining its circumscription so it remains monophyletic by including the *Pelagibacterales* order. The emended *Caulobacteridae* subclass within the *Alphaproteobacteria* class is defined based on phylogenetic analyses of 200 genes which are predominantly single-copy and vertically-inherited (unlikely laterally transferred) when compositionally heterogeneity was decreased by site removal or recoding. Phylogenetic (node-based) definition: the least inclusive clade containing *Pelagibacter* sp. HIMB058, ‘*Candidatus* Pelagibacter sp.’ IMCC9063, alphaproteobacterium HIMB59, *Caedibacter* sp. 37-49, ‘*Candidatus* Nucleicultrix amoebiphila’ FS5, ‘*Candidatus* Finniella lucida’, *Holospora obtusa* F1, *Sneathiella glossodoripedis* JCM 23214, *Sphingomonas wittichii*, and *Brevundimonas subvibrioides* ATCC 15264. The *Caulobacteridae* does not include: *Anaplasma phagocytophilum* HZ, *Rickettsia typhi* Wilmington, and ‘*Candidatus* Midichloria mitochondrii’ IricVA

*Azospirillaceae* fam. nov. (*Rhodospirillales*, *Alphaproteobacteria*) *Azospirillum* is the type genus of the family. This new family within the *Rhodospirillales* order is defined based on phylogenetic analyses of 200 genes which are predominantly single-copy and vertically-inherited (unlikely laterally transferred). Phylogenetic (node-based) definition: the least inclusive clade containing *Micavibrio aeruginoavorus* ARL-13, *Rhodocista centenaria* SW, and *Inquilinus limosus* DSM 16000. The *Azospirillaceae* does not include: *Rhodovibrio salinarum* DSM 9154, ‘*Candidatus* Puniceispirillum marinum’ IMCC 1322, *Rhodospirillum rubrum* ATCC 11170, *Terasakiella pusilla* DSM 6293, *Acidiphilium angustum* ATCC 49957, and *Elioraea tepidiphila* DSM 17972.

*Rhodovibriaceae* fam. nov. (*Rhodospirillales*, *Alphaproteobacteria*) *Rhodovibrio* is the type genus of the family. This new family within the *Rhodospirillales* order is defined based on phylogenetic analyses of 200 genes which are predominantly single-copy and vertically-inherited (unlikely laterally transferred). Phylogenetic (node-based) definition: the least inclusive clade containing *Rhodovibrio salinarum* DSM 9154, *Kiloniella laminariae* DSM 19542, *Oceanibaculum indicum* P24, *Thalassobaculum salexigens* DSM 19539 and ‘*Candidatus* Puniceispirillum marinum’ IMCC 1322. The *Rhodovobriaceae* does not include: *Rhodospirillum rubrum* ATCC 11170, *Terasakiella pusilla* DSM 6293, *Rhodocista centenaria* SW, *Micavibrio aeruginoavorus* ARL-13, *Acidiphilium angustum* ATCC 49957, and *Elioraea tepidiphila* DSM 17972.

*Rhodospirillaceae* emend. (*Rhodospirillales*, *Alphaproteobacteria*) *Rhodospirillum* is the type genus of the family. The *Rhodospirillaceae* family is here amended by redefining its circumscription so it remains monophyletic. The emended *Rhodospirillaceae* family within the *Rhodospirillales* order is defined based on phylogenetic analyses of 200 genes which are predominantly single-copy and vertically-inherited (unlikely laterally transferred). Phylogenetic (node-based) definition: the least inclusive clade containing *Rhodospirillum rubrum* ATCC 11170, *Roseospirillum parvum* 930l, *Magnetospirillum magneticum* AMB-1 and *Terasakiella pusilla* DSM 6293. The *Rhodospirillaceae* does not include: *Rhodocista centenaria* SW, *Micavibrio aeruginoavorus* ARL-13, ‘*Candidatus* Puniceispirillum marinum’ IMCC 1322, *Rhodovibrio salinarum* DSM 9154, *Elioraea tepidiphila* DSM 17972, and *Acidiphilium angustum* ATCC 49957.

Holosporaceae (*Rhodospirillales*, *Alphaproteobacteria*) *Holospora* is the type genus of the family. The *Holosporaceae* family as defined here has the same taxon circumscription as the *Holosporales* order *sensu* Szokoli et al., (2016), but it is here lowered to the family level and placed within the *Rhodospirillales* order. The new family rank-level for this group is based on the phylogenetic analysis of 200 genes, which are predominantly single-copy and vertically-inherited (unlikely laterally transferred), when compositionally heterogeneity was decreased by site removal or recoding (and coupled to the removal of the long-branching taxa *Pelagibacterales* and *Rickettsiales*). The family contains three subfamilies (lowered in rank from a former family level) and one formally-undescribed clade, namely, the *Holosporodeae*, and ‘*Candidatus Paracaedibacteriodeae’*, ‘*Candidatus Hepatincolodeae’*, and the *Caedibacter-Nucleicultrix* clade.

## METHODS

### Genome sequencing

Axenic cultures of *Viridiraptor invadens* strain Virl02, the host of ‘*Candidatus* Finniella inopinata’, were grown on the filamentous green alga *Zygnema pseudogedeanum* strain CCAC 0199 as described in (Hess and Melkonian 2013). Once the algal food was depleted, *Viridiraptor* cells were harvested by filtration through a cell strainer (mesh size 40 μm to remove algal cell walls) and centrifugation (~1,000 g for 15 min). For short-read sequencing, DNA extraction of total gDNA was carried out with the ZR Fungal/Bacterial DNA MicroPrep Kit (Zymo Research) using a BIO101/Savant FastPrep FP120 high-speed bead beater and 20 uL of proteinase K (20 mg/mL). A sequencing library was made using the NEBNext Ultra II DNA Library Prep Kit (New England Biolabs). DNA sequencing libraries were sequenced with an Illumina MiSeq instrument (Dalhousie University; Canada). For long-read sequencing, DNA extraction was performed using a CTAB and phenol-chloroform method. Total gDNA was further cleaned through a QIAGEN Genomic-Tip 20/G. A sequencing library was made using the Nanopore Ligation Sequencing Kit 1D (SQK-LSK108). Sequencing was done on a portable MinION instrument (Oxford Nanopore Technologies).

*Peranema trichophorum* strain CCAP 1260/1B was obtained from the Culture Collection of Algae and Protozoa (CCAP, Oban, Scotland) and grown in liquid Knop media plus egg yolk crystals. Total gDNA was extracted following Lang and Burger 2007. A sequencing library was made using a TruSeq DNA Library Prep Kit (Illumina). DNA sequencing libraries were sequenced with an Illumina MiSeq instrument (Genome Quebec Innovation Centre; Canada).

*Stachyamoeba lipophora* strain ATCC 50324 cells feeding on *Escherichia coli* were harvested and these were broken up with pestle and mortar in the presence of glass beads (< 450 μm diameter). Total gDNA was extracted using the QIAGEN Genomic G20 Kit. A sequencing library was made using a TruSeq DNA Library Prep Kit (Illumina). DNA sequencing libraries were sequenced with an Illumina MiSeq instrument (Genome Quebec Innovation Centre; Canada).

### Genome assembly and annotation

Short sequencing reads produced in an Illumina MiSeq from *Viridiraptor invadens*, *Peranema trichophorum*, and *Stachyamoeba lipophora* were first assessed with FASTQC v0.11.6 and then, based on its reports, trimmed with Trimmomatic v0.32 (Bolger, Lohse, and Usadel 2014) using the options: HEADCROP:16 LEADING:30 TRAILING:30 MINLEN:36. Illumina adapters were similarly removed with Trimmomatic v0.32 using the option ILLUMINACLIP. Long sequencing reads produced in an Nanopore MinION instrument from *Viridiraptor invadens* were basecalled with Albacore v2.1.7, adapters were removed with Porechop v0.2.3, lambda phage reads were removed with NanoLyse v0.5.1, quality filtering was done with NanoFilt v2.0.0 (with the options ‘--headcrop 50 -q 8 -l 1000’), and identity filtering against the high-quality short Illumina reads was done with Filtlong v0.2.0 (and the options ‘--keep_percent 90 --trim --split 500 --length_weight 10 --min_length 1000’). Statistics were calculated throughout the read processing workflow with NanoStat v0.8.1 and NanoPlot v1.9.1. A hybrid co-assembly of both processed Illumina short reads and Nanopore long reads from *Viridiraptor invadens* was done with SPAdes v3.6.2 (Bankevich et al. 2012). Assemblies of the Illumina short reads from *Peranema trichophorum* and *Stachyamobea lipophora* were separately done with SPAdes v3.6.2 (Bankevich et al. 2012). The resulting assemblies for both *Viridiraptor invadens* and *Peranema trichophorum* were later separately processed with the Anvi’o v2.4.0 pipeline (Eren et al. 2015) and refined genome bins corresponding to ‘*Candidatus* Finniella inopinata’ and the *Peranema*-associated rickettsialean were isolated primarily based on tetranucleotide sequence composition and taxonomic affiliation of its contigs. A single contig corresponding to the genome of the *Stachyamoeba*-associated rickettsialean was obtained from its assembly and this was circularized by collapsing the overlapping ends of the contig. Gene prediction and genome annotation was carried out with Prokka v.1.13 (see Table 1).

### Taxon and gene selection

The selection of 120 taxa was largely based on the phylogenetically diverse set of alphaproteobacteria determined by Wang and Wu (2015). To this set of taxa, recently sequenced and divergent unaffiliated alphaproteobacteria were added, as well as those claimed to constitute novel order-level taxa. Some other groups, like the *Pelagibacterales*, *Rhodospirillales* and the *Holosporales*, were expanded to better represent their diversity (see Fig. S1).

A set of 200 gene markers (54,400 sites; 9.03% missing data, see Fig. S1) defined by Phyla-AMPHORA was used (Wang and Wu 2013). The genes are single-copy and predominantly vertically-inherited as assessed by congruence among them (Wang and Wu 2013). Another smaller dataset of 40 compositionally-homogenous genes (5,570 sites; 5.98% missing data) was built by selecting the least compositionally heterogeneous genes from the larger 200 gene set according compositional homogeneity tests performed in P4 (Table S1). This was done as an alternative way to overcome the strong compositional heterogeneity observed in datasets for the *Alphaproteobacteria* with a broad selection of taxa. In brief, the P4 tests rely on simulations based on a provided tree (here inferred for each gene under the model LG4X+F in IQ-TREE) and a model (LG+F+G4 available in P4) to obtain proper null distributions to which to compare the *X*2 statistic. Most standard tests for compositional homogeneity (those that do not rely on simulate the data on a given tree) ignore correlation due to phylogenetic relatedness, and can suffer from a high probability of false negatives (Foster 2004).

Variations of our full set were made to specifically assess the placement of each long-branching groups individually. In other words, each group with comparatively long branches (the *Rickettsiales*, *Pelagibacterales*, *Holosporales*, and alphaproteobacterium sp. HIMB59) was analyzed in isolation, i.e., in the absence of other long-branching taxa. This was done with the purpose of removing the potential artefactual attraction among these groups. Taxon removal was done in addition to compositionally biased site removal and data recoding into reduced character-state alphabets (for a summary of the different methodological strategies employed see Fig. S2).

### Removal of compositionally biased and fast sites

As an effort to reduce artefacts in phylogenetic inference from our dataset (which might stem from extreme divergence in the evolution of the *Alphaproteobacteria*), we removed sites estimated to be highly compositionally heterogeneous or fast evolving. The compositional heterogeneity of a site was estimated by using a metric intended to measure the degree of disparity between the most %AT-rich taxa and all others. Taxa were ordered from lowest to highest proteome GARP:FIMNKY ratios; ‘GARP’ amino acids are encoded by %GC-rich codons, whereas ‘FIMNKY’ amino acids are encoded by %AT-rich codons. The resulting plot was visually inspected and a GARP:FIMNKY ratio cutoff of 1.06 (which represented a discontinuity or gap in the distribution) was chosen to divide the dataset into low GARP:FMINKY (or %AT-rich) and higher GARP:FIMNKY (or ‘GC-rich’) taxa (Fig. S3). Next, we determined the degree of compositional bias per site (𝒵) for the frequencies of both FIMNKY and GARP amino acids between the %AT-rich and all other (‘GC-rich’) alphaproteobacteria. To calculate this metric for each site the following formula was used:

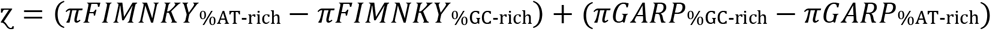

where *πFIMNKY* and *πGARP* are the sum of the frequencies for FIMNKY and GARP amino acids at a site, respectively, for either ‘%AT-rich’ or ‘%GC-rich’ taxa. According to this metric, higher values measure a greater disparity between %AT-rich alphaproteobacteria and all others; a measure of compositional heterogeneity or bias per site. The most compositionally heterogeneous sites according to 𝒵 were progressively removed using the software SiteStripper (Verbruggen 2018) in increments of 10%. We also progressively removed the fastest evolving sites in increments of 10%. Conditional mean site rates were estimated under the LG+C60+F+R6 model in IQ-TREE v1.5.5 using the ‘-wsr’ flag (Nguyen et al. 2015).

### Data recoding

Our datasets were recoded into four- and six-character state amino acid alphabets using dataset-specific recoding schemes aimed at minimizing compositional heterogeneity in the data (Susko and Roger 2007). The program minmax-chisq, which implements the methods of Susko and Roger (2007), was used to find the best recoding schemes—please see Fig. 3, S12 and S14 legends for the specific recoding schemes used for each dataset. The approach uses the chi-squared (*X*^2^) statistic for a test of homogeneity of frequencies as a criterion function for determining the best recoding schemes. Let *π_i_* denote the frequency of bin *i* for the recoding scheme currently under consideration. For instance, suppose the amino acids were recoded into four bins,

#### RNCM EHIPTWV ADQLKS GFY

Then *π*_4_ would be the frequency with which the amino acids G, F or Y were observed. Let *π_is_* be the frequency of bin *i* for the *s*th taxa. Then the *X*^2^ statistic for the null hypothesis that the frequencies are constant, over taxa, against the unrestricted hypothesis is

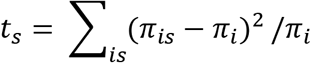

The *X*^2^ statistic provides a measure of how different the frequencies for the *s*th taxa are from the average frequencies. The maximum *t_s_* over *s* is taken as an overall measure of how heterogeneous the frequencies are for a given recoding scheme. The minmax-chisq program searches through recoding schemes, moving amino acids from one bin to another, to try to minimize the *max t_s_* (Susko and Roger 2007).

### Phylogenetic inference

The inference of phylogenies was primarily done under the maximum likelihood (ML) framework and using IQ-TREE v1.5.5 (Minh, Nguyen, and von Haeseler 2013; Nguyen et al. 2015). We first inferred guide trees (for a PMSF analysis) with a model that comprises the LG empirical matrix, with empirical frequencies estimated from the data (F), six rates for the FreeRate model to account for rate heterogeneity across sites (R6), and a mixture model with 60 amino acid profiles (C60) to account for compositional heterogeneity across sites—LG+C60+F+R6. Because the computational power and time required to properly explore the whole tree space (given such a big dataset and complex model) was too high, constrained tree searches were employed to obtain these initial guide trees (see Fig. S1 for the constraint tree). Many shallow nodes were constrained if they received maximum UFBoot and SH-aLRT support in a LG+PMSF(C60)+F+R6 analysis. All deep nodes, those relevant to the questions addressed here, were left unconstrained (Fig. S1). The guide trees were then used together with a dataset-specific mixture model ES60 to estimate site-specific amino acid profiles, or a PMSF (Posterior Mean Site Frequency Profiles) model, that best account for compositional heterogeneity across sites (Wang et al. 2018). The dataset-specific empirical mixture model ES60 also has 60 categories but, unlike the general C60, was directly estimated from our large dataset of 200 genes and 120 alphaproteobacteria using the methods described in (Susko, Lincker, and Roger 2018). Final trees were inferred using the LG+PMSF(ES60)+F+R6 model and a fully unconstrained tree search. Those datasets that produced the most novel topologies under ML were further analyzed under a Bayesian framework using PhyloBayes MPI v1.7 and the CAT-Poisson+Γ4 model (Lartillot and Philippe 2004; Lartillot, Lepage, and Blanquart 2009). This model allows for a very large number of classes to account for compositional heterogeneity across sites and, unlike in the more complex CAT-GTR+Γ4 model, also allows for convergence to be more easily achieved between MCMC chains. PhyloBayes MCMC chains were run for more than 10,000 cycles until convergence between the chains was achieved and the largest discrepancy (i.e., maxdiff parameter) was < 0.1. A consensus tree was generated from two PhyloBayes MCMC chains using a burn-in of 500 trees and sub-sampling every 10 trees.

Phylogenetic analyses of recoded datasets into four-character state alphabets were analyzed using IQ-TREE v1.5.5 and the model GTR+ES60S4+F+R6. ES60S4 is an adaptation of the dataset-specific empirical mixture model ES60 to four-character states. It is obtained by adding the frequencies of the amino acids that belong to each bin in the dataset-specific four-character state scheme S4 (see Data Recoding for details). Phylogenetics analyses of recoded datasets into six-character state alphabets were analyzed using PhyloBayes MPI v1.7 and the CAT-Poisson+Γ4 model. Maximum-likelihood analyses with a six-state recoding scheme could not be performed because IQ-TREE currently only supports amino acid datasets recoded into four-character states.

### Other analyses

The 16S rRNA genes of ‘*Candidatus* Finniella inopinata’, and the presumed endosymbionts of *Peranema trichophorum* and *Stachyamoeba lipophora* were identified with RNAmmer 1.2 server and BLAST searches. A set of 16S rRNA genes for diverse rickettsialeans and holosporaleans, and other alphaproteobacteria as outgroup, were retrieved from NCBI GenBank. The selection was based on Hess et al., (2016), Szokoli et al., (2016) and Wang and Wu (2015). Environmental sequences for uncultured and undescribed rickettsialeans were retrieved by keeping the 50 best hits resulting from a BLAST search of our three novel 16S rRNA genes against the NCBI GenBank non-redundant (nr) database. The sequences were aligned with the SILVA aligner SINA v1.2.11 and all-gap sites were later removed. Phylogenetic analyses on this alignment were performed on IQ-TREE v1.5.5 using the GTR+F+R8 model.

A UPGMA (average-linkage) clustering of amino acid compositions based on the 200 gene set for the *Alphaproteobacteria* was built in MEGA 7 (Kumar, Stecher, and Tamura 2016) from a matrix of Euclidean distances between amino acid compositions of sequences exported from the phylogenetic software P4 (http://p4.nhm.ac.uk/index.html).

### Data availability

The genome of ‘*Candidatus* Finniella inopinata’, endosymbiont of *Peranema trichophorum* strain CCAP 1260/1B and endosymbiont of *Stachyamoeba lipophora* strain ATCC 50324 were deposited in NCBI GenBank under the BioProject PRJNA501864. Raw sequencing reads were deposited on the NCBI SRA archive under the BioProject PRJNA501864. Multi-gene datasets as well as phylogenetic trees inferred in this study were deposited at Mendeley Data under the DOI: http://dx.doi.org/10.17632/75m68dxd83.1.

## ACKNOWLEDGEMENTS

Sergio A. Muñoz-Gómez is supported by a Killam Predoctoral Scholarship and a Nova Scotia Graduate Scholarship. This work was supported by Natural Sciences and Engineering Research (NSERC) Discovery Grants 2016-06792 to A.J.R, RGPIN/05754-2015 to C.H.S, RGPIN/05286-2014 to G.B., and RGPIN-2017-05411 to B.F.L. We thank Jon Jerlström Hultqvist and Gina Filloramo (both at Dalhousie University) for advice on long-read sequencing with the Nanopore MinION. We also thank Camilo A. Calderón-Acevedo for advice about taxonomic issues, and Franziska Szokoli for reading and commenting on a late version of this manuscript. Bruce Curtis (Dalhousie University) and Peter G. Foster (Natural History Museum of London) kindly provided technical help with bioinformatics and with the software P4, respectively. We thank Joanny Roy, Georgette Kiethega, Matus Valach, and Shona Teijeiro (all at the Université de Montréal), and Drahomira Faktora (University of South Bohemia), for help with the culturing, DNA preparation and sequencing of the endosymbiont of *Peranema trichophorum*. Some of the genome data used in this study were produced by the US Department of Energy Joint Genome Institute (http://www.jgi.doe.gov/) in collaboration with the user community.

## SUPPLEMENTARY FIGURE LEGENDS

**Figure S1.** Constraint tree, used for IQ-TREE analyses, labeled with taxon names and also degree of missing data per taxon.

**Figure S2.** A diagram of the strategies employed in this study.

**Figure S3.** GARP:FIMNKY ratios across the proteomes of the 120 alphaproteobacteria used in this study.

**Figure S4.** A 16S rRNA gene maximum-likelihood tree of the *Rickettsiales* and *Holosporales* that phylogenetically places the three endosymbionts whose genomes were sequenced in this study: (1) ‘*Candidatus* Finniella inopinata’ endosymbiont of *Viridiraptor invadens* strain Virl02, (2) an alphaproteobacterium associated with *Peranema trichophorum* strain CCAP 1260/1B, and (3) an alphaproteobacterium associated with *Stachyamoeba lipophora* strain ATCC 50324. Branch support values are SH-aLRT and UFBoot.

**Figure S5.** A labeled version of Figure 2. Branch support values are 100% SH-aLRT and 100% UFBoot unless annotated.

**Figure S6.** Bayesian consensus trees inferred with PhyloBayes MPI v1.7 and the CAT-Poisson+Γ4 model. Branch support values are 1.0 posterior probabilities unless annotated. **A.** Bayesian consensus phylogram inferred from the full dataset which is highly compositionally heterogeneous. **B.** Bayesian consensus phylogram inferred from a dataset whose compositional heterogeneity has been decreased by removing 50% of the most biased sites according to 𝒵. See Figs. 2A and 2B for the most likely trees inferred in IQ-TREE v1.5.5 and the LG+PMSF(C60)+F+R6 model.

**Figure S7.** Ultrafast bootstrap (UFBoot) variation as the fastest sites are progressively removed in steps of 10%.

**Figure S8.** Ultrafast bootstrap (UFBoot) variation as compositionally biased sites, according to 𝒵, are progressively removed in steps of 10%.

**Figure S9.** Bayesian consensus trees inferred with PhyloBayes MPI v1.7 and the CAT-Poisson+Γ4 model. Branch support values are 1.0 posterior probabilities unless annotated. **A.** Bayesian consensus phylogram inferred to place the *Holosporales* in the absence of the *Rickettsiales* and the *Pelagibacterales* and when compositional heterogeneity has been decreased by removing 50% of the most biased sites according to 𝒵. **B.** Bayesian consensus phylogram inferred to place the *Holosporales* in the absence of the *Rickettsiales* and the *Pelagibacterales* and when the data have been recoded into a four-character state alphabet (the dataset-specific recoding scheme S4: ARNDQEILKSTV GHY CMFP W) to reduce compositional heterogeneity. See Figs. 2A and 2B for the most likely trees inferred in IQ-TREE v1.5.5 and the LG+PMSF(C60)+F+R6 and GTR+ES60S4+F+R6 models, respectively.

**Figure S10.** Maximum-likelihood phylograms obtained from the full dataset and after removing long-branching taxa. **A.** A phylogram for which all long-branching taxa are included. **B.** A phylogram to assess the placement of the *Holosporales*. **C.** A phylogram to assess the placement of the *Rickettsiales*. **D.** A phylogram to assess the placement of the *Pelagibacterales*. **E.** A phylogram to assess the placement of alphaproteobacterium sp. HIMB59.

**Figure S11.** Maximum-likelihood phylograms obtained after removing 50% of the most compositionally biased sites and removing long-branching taxa. **A.** A phylogram for which all long-branching taxa are included. **B.** A phylogram to assess the placement of the *Holosporales*. **C.** A phylogram to assess the placement of the *Rickettsiales*. **D.** A phylogram to assess the placement of the *Pelagibacterales*. **E.** A phylogram to assess the placement of alphaproteobacterium sp. HIMB59.

**Figure S12.** Maximum-likelihood phylograms obtained after recoding the data into S4 and removing long-branching taxa. **A.** A phylogram for which all long-branching taxa are included (recoding scheme: RNCM EHIPTWV ADQLKS GFY). **B.** A phylogram to assess the placement of the *Holosporales* (recoding scheme: ARNDQEILKSTV GHY CMFP W). **C.** A phylogram to assess the placement of the *Rickettsiales* (recoding scheme: PY RNMF GHLKTW ADCQEISV). **D.** A phylogram to assess the placement of the *Pelagibacterales*. **E.** A phylogram to assess the placement of alphaproteobacterium sp. HIMB59 (recoding scheme: RLKMT ANDQEIPSV CW GHFY).

**Figure S13.** Maximum-likelihood phylograms obtained from the 40 most compositionally homogeneous genes and removing long-branching taxa. **A.** A phylogram for which all long-branching taxa are included. **B.** A phylogram to assess the placement of the *Holosporales*. **C.** A phylogram to assess the placement of the *Rickettsiales*. **D.** A phylogram to assess the placement of the *Pelagibacterales*. **E.** A phylogram to assess the placement of alphaproteobacterium sp. HIMB59.

**Figure S14.** Bayesian consensus phylogram inferred to place the *Holosporales* in the absence of the *Rickettsiales* and the *Pelagibacterales* and when the data have been recoded into a six-character state alphabet (the dataset-specific recoding scheme S6: AQEHISV RKMT PY DCLF NG W) to reduce compositional heterogeneity. Branch support values are 1.0 posterior probabilities unless annotated.

**Table S1.** A list of the least compositionally heterogeneous genes out of the 200 single-copy and vertically-inherited genes used in this study.

